# Potent antibody immunity to SARS-CoV-2 variants elicited by a third dose of inactivated vaccine

**DOI:** 10.1101/2021.11.10.468037

**Authors:** Bin Ju, Bing Zhou, Shuo Song, Qing Fan, Xiangyang Ge, Haiyan Wang, Lin Cheng, Huimin Guo, Dan Shu, Lei Liu, Zheng Zhang

**Affiliations:** Institute for Hepatology, National Clinical Research Center for Infectious Disease, Shenzhen Third People’s Hospital; The Second Affiliated Hospital, School of Medicine, Southern University of Science and Technology, Shenzhen 518112, Guangdong Province, China; Shenzhen Research Center for Communicable Disease Diagnosis and Treatment of Chinese Academy of Medical Science, Shenzhen 518112, Guangdong Province, China; Guangdong Key laboratory for anti-infection Drug Quality Evaluation, Shenzhen 518112, Guangdong Province, China; Department for Infectious Diseases, National Clinical Research Center for Infectious Disease, Shenzhen Third People’s Hospital; The Second Affiliated Hospital, School of Medicine, Southern University of Science and Technology, Shenzhen 518112, Guangdong Province, China

**Author notes:** These authors contributed equally to this work (Bin Ju, Bing Zhou, Shuo Song). Correspondence (Z.Z.), (L.L.), (B.J.).

## Abstract

SARS-CoV-2 variants are still prevalent worldwide and continue to pose a challenge to the effectiveness of current vaccines. It remains unknown whether a third dose of inactivated vaccine elicits immune potential against SARS-CoV-2 variants. Here, we showed a significant decline in plasma neutralization against SARS-CoV-2 at seven months after a second dose of the inactivated vaccine in a large-scale cohort. However, we also found that a third vaccination with an inactivated vaccine largely increased plasma neutralization against variants including Beta, Delta, and Lambda. More importantly, the high-affinity anti-RBD memory B cells were also generated by the third vaccination, suggesting a more potent and longer protection. These findings highlighted the importance and effectiveness of a third dose of inactivated vaccine in conferring higher protection against the emerging variants in populations.

## Introduction

The coronavirus disease 2019 (COVID-19) pandemic has already lasted for nearly two years and continues to threaten human health and life. By October 26, 2021, severe acute respiratory syndrome coronavirus 2 (SARS-CoV-2) had infected more than 244 million individuals and caused over 4.9 million deaths around the world. While several effective vaccines have been deployed to combat wild-type (WT) virus infection^1-3^, the emerging SARS-CoV-2 variants showed enhanced transmissibility and significantly escaped the neutralization of vaccine-elicited plasma. New infections caused by SARS-CoV-2 variants are still rising sharply worldwide, especially the Alpha, Beta, Delta, and Lambda variants. They have contributed to several current waves of infection globally^4-9^.

More seriously, breakthrough infections of SARS-CoV-2 variants after vaccination have occurred widely with a significant reduction in vaccine efficacy over time^10-12^. Data from a recent study in New York City demonstrated that mutated strains, including Alpha and Iota, are able to escape the protection of several vaccines, including BNT162b2, mRNA-1273 and JNJ-78436735^13^. The remarkable drop in neutralizing activities of vaccine-elicited plasma has been considered to be a key factor leading to breakthrough infection.

Currently, many researchers have asked whether a third dose of vaccine is necessary to increase the titers of neutralizing antibodies (nAbs) against SARS-CoV-2 variants and to better control the current COVID-19 pandemic. A third dose of the mRNA vaccine BNT162b2 has been proven to be effective in combatting variants. It was reported that the neutralization geometric mean titers (GMTs) against Beta increased more than 15 to 20 times compared with those after the second vaccination, and the ratio (Delta to WT) of neutralization GMTs raised to 0.85 and 0.92 in younger adults and in older adults after a third dose of BNT162b2, respectively^14^.

The inactivated vaccine, as an important vaccine candidate, has shown good immunogenicity in clinical trials and has been widely used in the population^15,16^. However, it remains elusive whether plasma antibody titers against SARS-CoV-2 variants decline with time in inactivated vaccinees, especially in those who have received two doses of vaccines for more than half a year. In addition, the antibody immunity to SARS-CoV-2 variants elicited by a third dose of inactivated vaccine has not been comprehensively analyzed in a large-scale cohort, which is critical to develop strategies for curbing the spread of SARS-CoV-2 variants.

In this study, we summarized a large cohort of more than 500 individuals who received two or three doses of inactivated SARS-CoV-2 vaccines (BBIBP-CorV) and were followed up for nearly nine months. We characterized the kinetics of plasma IgG and IgM bound to the viral receptor binding domain (RBD) throughout the follow-up period and defined the decline in neutralizing activities against SARS-CoV-2 Beta, Delta, and Lambda variants in inactivated vaccinees 7 months after the second vaccination. More importantly, we proved that a third dose of inactivated vaccine could significantly increase the titers of binding and neutralizing antibodies against SARS-CoV-2 variants and enhance the percentages and affinities of RBD-specific memory B cells (MBCs). These data provided a proof of concept that a third booster immunization with an inactivated vaccine could be considered an effective measure against the SARS-CoV-2 variant pandemic.

## Methods

### Study approval and blood samples

This study was approved by the Ethics Committee of Shenzhen Third People’s Hospital, China (approval number: 2020-030). All participants had provided written informed consent for sample collection and subsequent analysis. All plasma and peripheral blood mononuclear cells (PBMCs) from individuals who received two or three doses of inactivated SARS-CoV-2 vaccines (BBIBP-CorV, the Sinopharm COVID-19 vaccine, Beijing Institute of Biological Products Co., Ltd) were collected at different time points of follow-up from the Biobank of the Shenzhen Third People’s Hospital. All plasma samples were stored at -80 °C and heat-inactivated at 56 °C for 1 h before use. PBMCs were maintained in freezing medium and stored in liquid nitrogen.

### Enzyme linked immunosorbent assay (ELISA)

SARS-CoV-2 wild-type (WT) and mutated (Beta: K417N-E484K-N501Y, Delta: L452R-T478K, Lambda: L452Q-F490S) RBD proteins (Sino Biological) were separately coated into 96-well plates at 4 °C overnight. The plates were washed with PBST buffer and blocked with 5% skim milk and 2% bovine albumin in PBS at room temperature (RT) for 1 h. Plasma samples were diluted at 1:20, added to the wells, and then incubated at 37 °C for 1 h. The plates were washed, and HRP-conjugated goat anti-human IgG antibodies (ZSGB-BIO) were added and then incubated at 37 °C for 30 mins. Finally, the TMB substrate (Sangon Biotech) was added to the wells and incubated at RT for 5 mins, and the reaction was stopped with 2 M H_2_SO_4_. The readout was detected at wavelengths of 450 nm and 630 nm. For titration of the end-point titers of binding antibodies, plasma samples were serially diluted 3-fold from 1:20 to 1:43740 and then added to the plates. The following steps were the same as those mentioned above. A cutoff was set as an OD_450nm-630nm_ value of 0.100. The end-point titer was defined as the last dilution whose OD_450nm-630nm_ value was over 0.100.

### SARS-CoV-2 pseudovirus-based neutralizing assay

SARS-CoV-2 pseudovirus was generated by cotransfection of HEK-293T cells with SARS-CoV-2 spike-expressing plasmid and an env-deficient HIV-1 backbone vector (pNL4-3.Luc.R-E-). Two days post transfection, the culture supernatant was harvested, clarified by centrifugation, filtered and stored at - 80 °C. To determine the neutralizing activity, plasma samples were serially diluted and incubated with an equal volume of SARS-CoV-2 pseudovirus at 37 °C for 1 h. HEK-293T-hACE2 cells were subsequently added to the plates. After a 48 h incubation, the culture medium was removed, and 100 μL of Bright-Lite Luciferase reagent (Vazyme Biotech) was added to the cells. After a 2 min incubation at RT, 90 μl of cell lysate was transferred to 96-well white solid plates for measurements of luminescence using the Varioskan™ LUX multimode microplate reader (Thermo Fisher Scientific). The 50% inhibitory dilution (ID_50_) was calculated using GraphPad Prism 8.0 software by log (inhibitor) vs. normalized response - Variable slope (four parameters) model.

### Flow cytometric analysis of RBD-specific memory B cells

Thawed PBMCs were stained with an antibody cocktail consisting of CD19- PE-Cy7, CD3-Pacific Blue, CD8-Pacific Blue, CD14-Pacific Blue, CD27-APC-H7, and IgG-FITC (all from BD Biosciences) to gate IgG^+^ memory B cells. SARS-CoV-2 WT RBD with His tag (Sino Biological) was used as a probe to target antigen-specific B cells. Two anti-His secondary antibodies separately labeled with APC and PE (Abcam) were both used to recognize the RBD bait and exclude nonspecific staining. A LIVE/DEAD Fixable Dead Cell Stain Kit (Invitrogen) was used to exclude dead cells. Flow cytometric data were acquired on an Aria II flow cytometer (BD Biosciences) and analyzed using FlowJo software (TreeStar).

### Statistical analysis

Statistical analysis was performed with paired or unpaired t tests using GraphPad Prism 8.0 software. *, P < 0.05; **, P < 0.01; ***, P < 0.001; ****, P < 0.0001.

## Results

### Longitudinal dynamics of plasma IgG and IgM against SARS-CoV-2 during three doses of inactivated vaccines

Five hundred and thirty-three participants who received two or three doses of BBIBP-CorV containing 4 μg total protein were enrolled in this study. These donors were followed up at Week 2 after first vaccination (n = 344), Week 2 after second vaccination (n = 533), Month 2 after second vaccination (n = 286), Month 7 after second vaccination (i.e., before third vaccination, n = 130), and Week 2 after third vaccination (n = 176). As shown in Figure 1A, a total of 1469 blood samples were collected from 533 donors at the above five follow-up time points.

**Figure 1.**
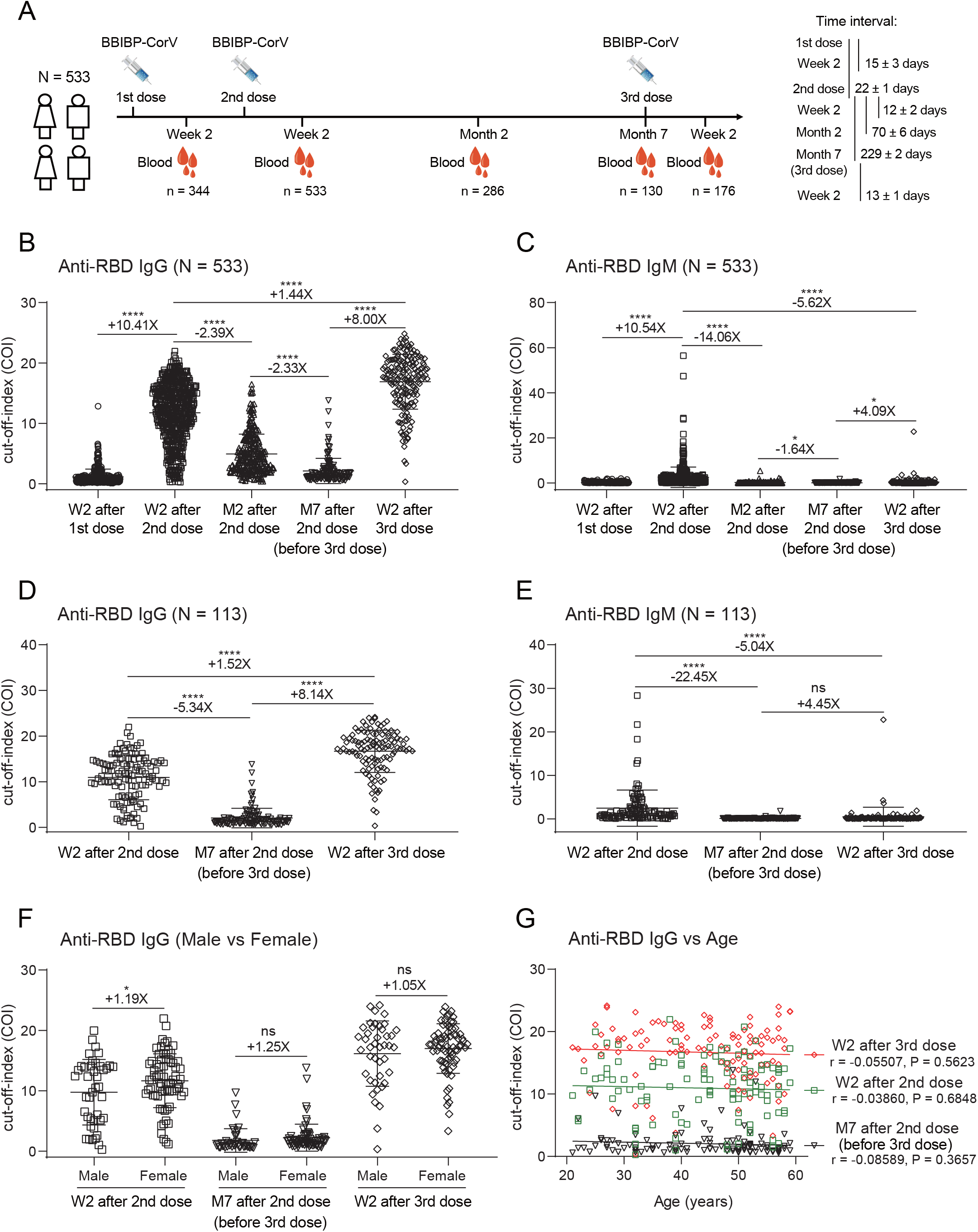
Longitudinal dynamics of humoral antibodies and boosting effect of a third dose of inactivated vaccine against SARS-CoV-2. **(A)** Immunization schedule and blood specimen collection of 533 donors who received two or three doses of inactivated vaccines in this project. The interval time is shown as the mean ± SD days. **(B-C)** Plasma antibody dynamics of anti-RBD IgG **(B)** and IgM **(C)** during three doses of vaccines. **(D-E)** The binding ability of IgG **(D)** and IgM **(E)** to RBD from 113 donors who completed three time-point follow-up visits: Week 2 and Month 7 after second vaccination and Week 2 after third vaccination. **(F)** Comparison of anti-RBD IgG values between male (n = 42) and female (n = 71) vaccinees. **(G)** Correlation analysis between anti-RBD IgG values and ages in the 113 donors at three follow-up visits.

We measured the binding IgG and IgM activities to SARS-CoV-2 WT RBD in all plasma using a chemiluminescence immunoassay kit. IgG seroconversion was present in more than 98% (526/533) of vaccine recipients at Week 2 after second vaccination. The mean value of plasma IgG was significantly increased 10.41-fold compared with that at Week 2 after first vaccination. However, anti-RBD IgG values were gradually decreased to 41.8% at Month 2 after second vaccination and additionally dropped to 42.9% at Month 7. Specially, the positive rate of RBD-specific IgG was decreased to approximately 75% in 130 donors at Month 7 after second vaccination (Figure 1B).

Subsequently, we collected blood samples from a total of 176 individuals who accepted a third dose of BBIBP-CorV. Nearly all vaccinees (175/176) were characterized by the seroconversion of IgG against SARS-CoV-2 at Week 2 after third vaccination, and their mean IgG values were increased 8.00-fold compared with those before third vaccination and 1.44-fold compared with those after second vaccination (Figure 1B). RBD-specific IgM displayed a similar pattern of kinetics as IgG, although the mean values of IgM were absolutely lower than IgG at each time point (Figure 1C). However, the difference between IgM and IgG was that the third dose of vaccine did not induce a strong IgM response, suggesting that IgG may play a more important role in recalling to the SARS-CoV-2 vaccine or viral infection exposure.

### A third dose of inactivated vaccine elicits robust binding antibodies to SARS-CoV-2 independent on gender and age

To evaluate the effects of the third dose on humoral immune responses, we detected serially paired plasma samples before third vaccination (at Week 2 and Month 7 after second vaccination) and Week 2 after third vaccination in 113 donors. Seven months after second vaccination, RBD-specific IgG levels induced by two doses of vaccines were sharply decreased 82.1% compared with those at Week 2 after second vaccination. Importantly, the mean COI value of plasma anti-RBD IgG rapidly increased to 16.7 by a third vaccination, which was an 8.14-fold increase compared to that before third vaccination and was also significantly higher than that induced by two doses of vaccines (Figure 1D). In contrast, a third dose of inactivated vaccine failed to induce a recalling IgM response (Figure 1E).

We further compared the differences in anti-RBD IgG levels between male and female donors. There were 42 male donors (37%) and 71 female donors (63%) in the cohort. As shown in Figure 1F, both male and female donors displayed similar levels of RBD-specific IgG after third vaccination, although female donors had higher levels of anti-RBD IgG than males at Week 2 after second vaccination. There were no obvious relationships between anti-RBD IgG and ages at the three different follow-up time points (Figure 1G). These data showed that the third vaccination with an inactivated vaccine elicits robust binding antibodies to SARS-CoV-2 independent on gender and age.

### A third dose of inactivated vaccine elicits potent neutralizing antibodies against SARS-CoV-2 variants

To evaluate the ability of a third dose of inactivated vaccine to fight against the infection of mutant viruses, we detected both binding antibodies to RBD proteins and neutralizing antibodies (nAbs) against pseudoviruses of SARS-CoV-2 WT, Beta, Delta, and Lambda variants. We established the ELISA and pseudovirus-based neutralizing assay to test the binding activity and neutralization. The results of 20 non-vaccinated healthy donor plasma samples and a positive antibody control showed low backgrounds and good specificities of the two assays (Figure S1).

Then, we applied these two assays to detect the binding and neutralizing activities of plasma from the 113 donors. As shown in Figure 2A, similar to the binding response to WT RBD, mutated RBD-specific IgG was sharply decreased 7 months after second vaccination but was significantly increased by the third vaccination. The plasma neutralizing activities against WT and mutant pseudoviruses elicited by the inactivated vaccine displayed the same patterns as their binding activities (Figure 2B).

**Figure 2.**
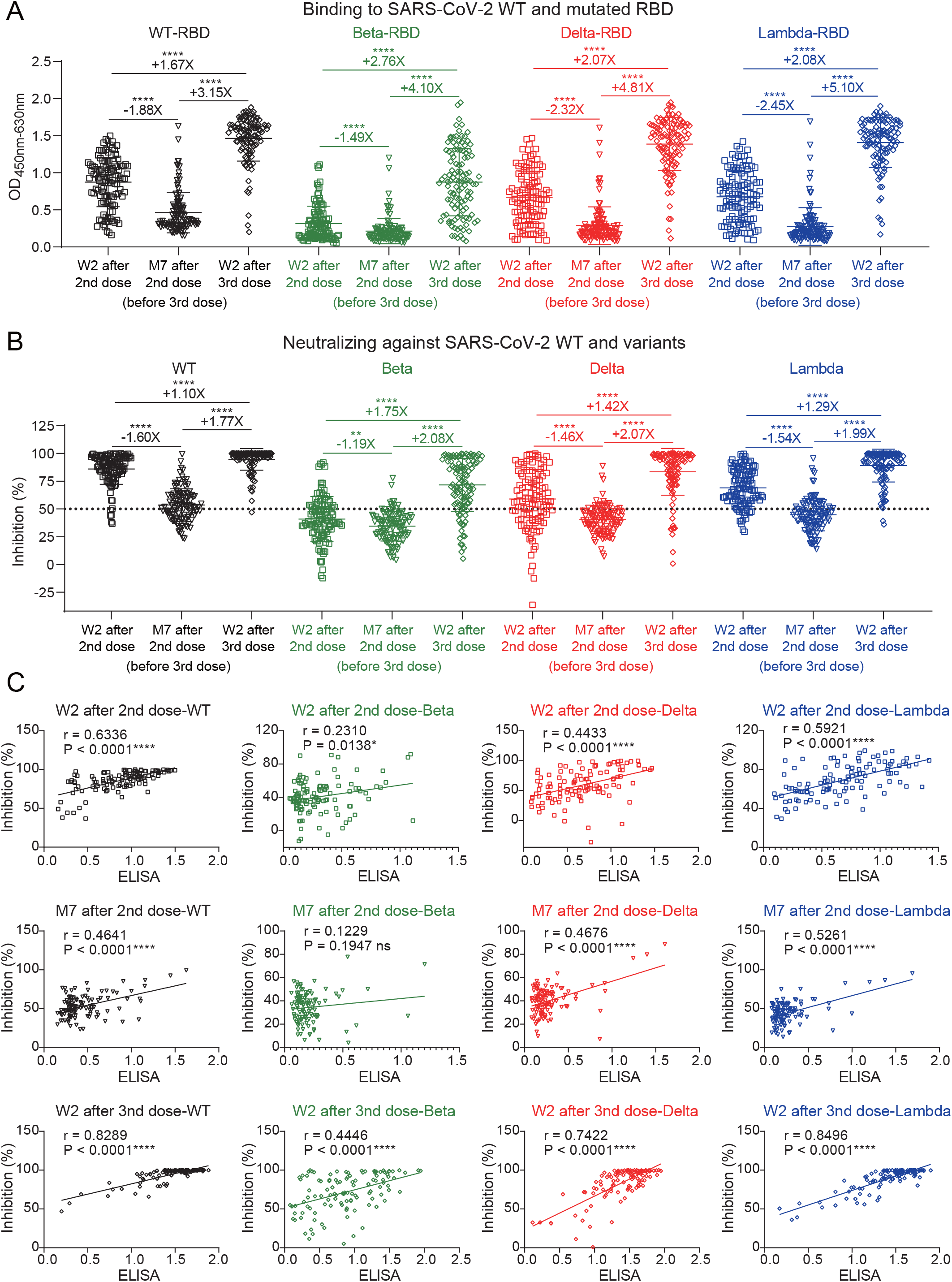
Potent binding and neutralizing antibodies against SARS-CoV-2 variants induced by a third dose of inactivated vaccine. **(A)** ELISA binding of 113 donors at three follow-up visits to SARS-CoV-2 WT and mutated RBD proteins. **(B)** Neutralizing activities of 113 donors at three follow-up visits against SARS-CoV-2 WT, Beta, Delta, and Lambda variants. The inhibition of 50% is indicated by a horizontal dashed line. **(C)** Correlation analysis between binding and neutralizing activities against SARS-CoV-2 WT and variants of 113 donors at three follow-up visits.

At Week 2 after second vaccination, 95% (108/113) of plasma demonstrated effective neutralization against WT virus with more than 50% inhibition at a 1:20 dilution. At this time point, they, to some extent, maintained neutralizing activities against some important SARS-CoV-2 variants (Beta, Delta, and Lambda). However, at Month 7 after second vaccination, nearly half of the plasma (49/113) lost their neutralizing activities (inhibition < 50%), and the inhibition was decreased to 53.6% against the WT strain in these 113 donors. Notably, the inhibitions of plasma against Beta, Delta and Lambda variants had decreased to 34.4%, 40.4%, and 44.8%, respectively, indicating their poor defenses against SARS-CoV-2 variants.

After the third vaccination, the plasma inhibitions against WT, Beta, Delta, and Lambda were significantly increased to 94.6%, 71.6%, 83.4%, and 89.0% within 2 weeks, respectively. The binding and neutralizing activities of plasma against these variants were strongly related to those against WT virus (Figure S2 and S3). In addition, there were significant correlations between plasma inhibitions and their binding activities at various time points after vaccination (Figure 2C). These findings indicated that a third dose of inactivated vaccine elicited potent neutralizing antibodies against SARS-CoV-2 variants.

### A third dose of inactivated vaccine elicits high-affinity memory B cells

Virus-specific MBCs play important roles in recalling to viral infection and partially contribute to the durability of antibody immunity. Therefore, we randomly selected 24 individuals and summarized their 72 PBMCs before third vaccination (at Week 2 and Month 7 after second vaccination) and Week 2 after third vaccination to detect SARS-CoV-2 RBD-specific MBCs (Figure S4). As shown in Figure 3A, although plasma anti-RBD IgG was gradually decreased over time, the percentage of RBD-specific MBCs maintained a similar level at Month 7 as that at Week 2 after second vaccination, suggesting that these vaccine recipients might still retain a certain protection against SARS-CoV-2 infection. Notably, the percentage of RBD-specific MBCs rapidly increased after the third vaccination, which was significantly higher than that at Week 2 after second vaccination and before third vaccination (0.96% vs. 0.50% and 0.53%). The mean fluorescence intensity (MFI) of RBD-binding MBCs was also significantly enhanced by the third vaccination compared to that at Week 2 after second vaccination and before third vaccination (4799 vs. 2951 and 2680 in APC, 8894 vs. 4516 and 4352 in PE). These data indicated that the third vaccination not only increases the proportion of MBCs but also enhances the RBD affinity with MBCs.

**Figure 3.**
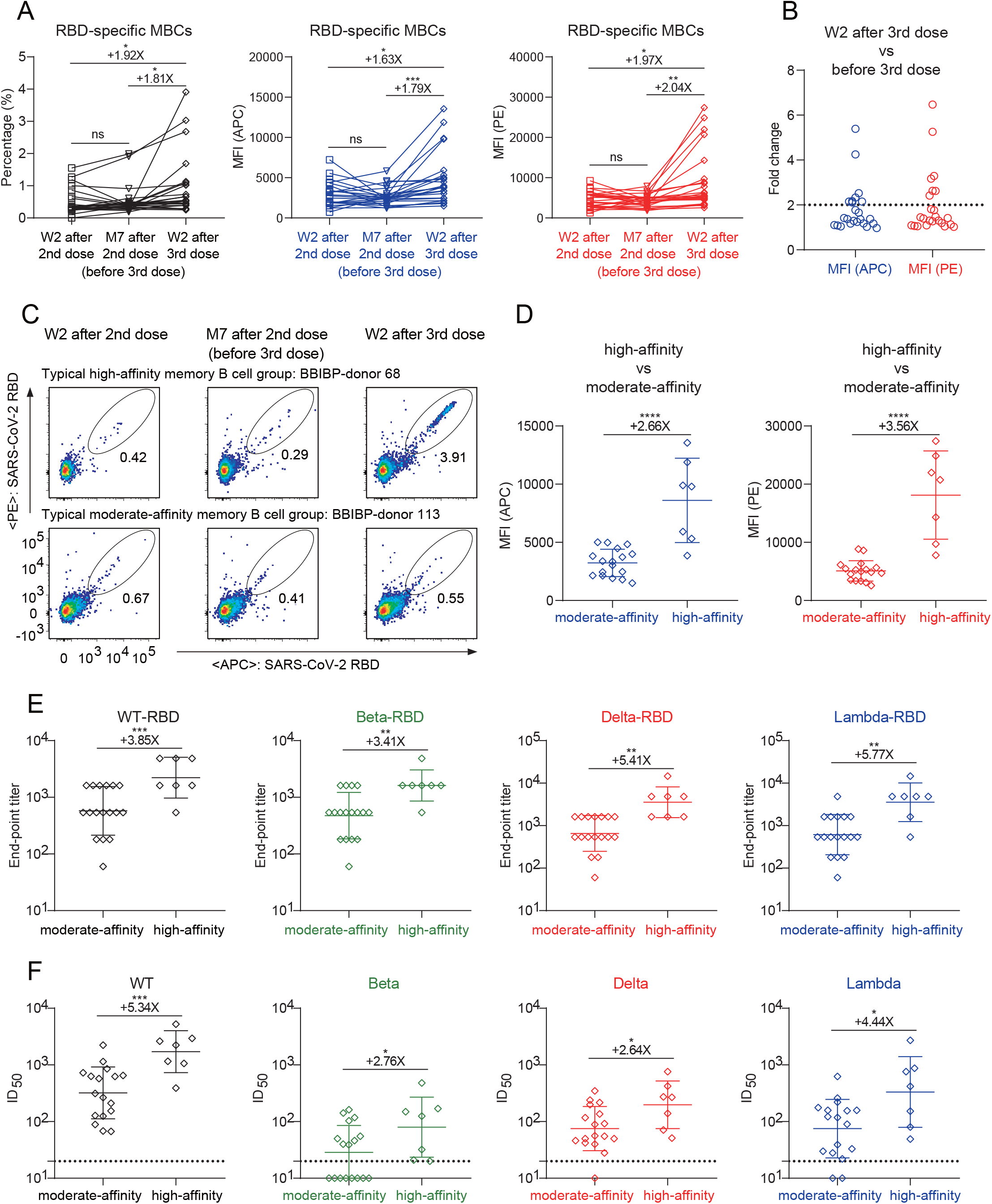
High-affinity RBD-specific memory B cells elicited by a third dose of inactivated vaccine. **(A)** The percentage (left), MFI of APC (middle), and MFI of PE (right) of RBD-specific MBCs (CD19^+^CD3^-^CD8^-^CD14^-^CD27^+^IgG^+^SARS-CoV-2-RBD^+^ cells) of randomly selected 24 donors with three follow-up visits. **(B)** The fold change of MFI in both APC and PE of RBD-specific MBCs between Week 2 after third vaccination and before third vaccination. A cutoff of 2-fold is indicated by the horizontal dashed line. High-affinity group: fold change > 2, moderate-affinity group: fold change < 2. **(C)** The typical display of high-affinity and moderate-affinity RBD-specific MBCs of 2 donors with three follow-up visits (high: BBIBP-donor 68, moderate: BBIBP-donor 113). **(D)** Comparison of MFI in both APC (left) and PE (right) of RBD-specific MBCs at Week 2 after third vaccination between the high-affinity group (n = 7) and the moderate-affinity group (n = 17). **(E)** The end-point titers of binding IgG to SARS-CoV-2 WT, Beta, Delta, and Lambda RBD proteins at Week 2 after third vaccination in the high-affinity and moderate-affinity groups. **(F)** The geometric mean titers of nAbs against SARS-CoV-2 WT, Beta, Delta, and Lambda pseudoviruses at Week 2 after third vaccination in the high-affinity and moderate-affinity groups. A cutoff of 1:20 dilution is indicated by a horizontal dashed line.

More importantly, we found that the third dose of vaccine induced extremely high-affinity MBCs in some individuals, whose MFIs of APC and PE on RBD-binding MBCs were both more than 2-fold higher than before the third vaccination (Figure 3B and 3C). Seven individuals with more than 2-fold higher MFI were defined as the high-affinity group, while the other 17 individuals were defined as the moderate-affinity group (Figure 3D). We thus compared the binding and neutralizing activities of plasma between the high-affinity and moderate-affinity groups at Week 2 after third vaccination. The geometric mean end-point titers of the plasma binding antibodies were significantly higher in the high-affinity group than those in the moderate-affinity group against WT RBD and the other mutated RBD proteins (Beta, Delta, and Lambda) (Figure 3E and S5). Similarly, the neutralizing activities of plasma were also significantly higher in the high-affinity group than those in the moderate-affinity group (Figure 3F and S6). Therefore, the third vaccination of inactivated vaccine indeed elicited more potent nAbs and high-affinity MBCs, suggesting a persistent antibody immunity to SARS-CoV-2 variants.

## Discussion

A large number of nAbs recognizing the RBD of virus have been isolated and divided into four classes according to their competitions with cell receptor (ACE2) and accessibilities of binding epitopes on the RBD in ‘up’ or ‘down’ conformations^17^. These anti-RBD nAbs could totally destroy or partly disturb the RBD-ACE2 interaction and thus block virus entry effectively. However, SARS-CoV-2 variants, including emerging variants of concern (VOCs) and variants of interest (VOIs), carry various mutations in the region of spike, especially on the nAb-binding sites of RBD. These mutations located in or near recognizing epitopes may lead to a significant decline in the neutralization of nAbs^5,6,18,19^.

One of the early VOCs, Beta, was first reported in South Africa and had the greatest reduction in neutralization capacity thus far^20^. Delta was identified in India and rapidly spread to many other countries, which had been classified as another VOC with 60% more transmissibility than Alpha and led to the current wave of COVID-19 pandemic^7^. In addition, recent preprint papers reported that a new VOI-Lambda with various deletions and substitutions in spike exhibited high infectivity and antibody resistance^9,21^.

Current vaccines are derived from the original Wuhan-Hu-1 gene, many nAbs elicited by which have been escaped due to the viral mutation. The K417N and E484K substitutions in Beta severely disrupted the binding of Class 1 and Class 2 nAbs to the RBD^5,22^. The L452R/Q mutant led to Delta and Lambda escaping from the neutralization of nAbs from Class 3^7,9,21^. Encouragingly, some nAbs, such as Class 4, still neutralize the above variants effectively^23-25^, explaining why the plasma of vaccine recipients and convalescent individuals maintained on some extent neutralizing activities. These residual broad nAbs play important roles in fighting against SARS-CoV-2 variants, leading to develop several strategies to increase their antibody titers. Booster immunization with the original vaccine is usually regarded as a direct and effective way to rapidly enhance the antibody titer and defend against the variants. Several evidences have demonstrated that vaccine recipients boosted with another dose of viral vector-based vaccine or mRNA vaccine rapidly produced sufficient nAbs against variants including Alpha, Beta, Gamma, and Delta^14,26^.

It remains unknown whether a third vaccination with an inactivated vaccine induces protective antibody responses against SARS-CoV-2 variants. We therefore evaluated the kinetics of plasma nAbs against variants and RBD-specific MBCs in a large-scale cohort who received two or three doses of inactivated vaccines. Although plasma neutralizing activity was generally reduced at 7 months after second vaccination, the antibody memory responses were well established by two doses of inactivated vaccines. The third dose of inactivated vaccine rapidly and significantly increased plasma antibody titers against various variants and generated high-affinity MBCs binding to RBD within two weeks. These findings suggest that a booster dose of inactivated vaccine increases the magnitude and breadth of neutralization in the pre-existing antibody response.

It is notable the differential dynamic of plasma neutralization and memory B cell responses by vaccination. Plasma neutralization peaks at two weeks post second vaccination and drops largely at seven months post second vaccination, then rebounds after the third vaccination. In contrast, MBCs are maintained at stable levels until seven months after second vaccination and are then significantly increased by a third vaccination. The mechanisms underlying the differential kinetics of plasma neutralization and memory B cell responses induced by vaccination are unclear. One possibility is that plasma neutralization, to a greater extent, reflects the functionality of the long-lived plasma cells in the bone marrow^27^. The MBC response is another important type of immune protection, whose quantity and quality contribute to the speed and potency of the immune system responding to viral reinfection^28^. Both factors collectively provide vaccine recipients with antibody protection against viral infection or prevent them from developing severe disease. Therefore, long-term monitoring of plasma neutralization against SARS-CoV-2 variants and RBD-specific MBCs is valuable for evaluating the vaccine effectiveness. Long-term follow-up will evaluate the duration of the antibody response elicited by the third dose of inactivated vaccine in future studies.

Overall, our data highlighted the challenges for vaccine recipients who have received complete immunization more than 6 months. They are at risk of breakthrough infection by SARS-CoV-2 variants, as plasma neutralization is generally reduced at half a year after the second vaccination. Meanwhile, viral evolution will continue and new variants will emerge one after another. Based on a large-scale cohort with a long follow-up time, we emphasize the importance of a third dose of SARS-CoV-2 inactivated vaccine to confer higher protection against emerging variants.

## Acknowledgments

We thank all participants who received inactivated vaccine and all of the healthy providers from Shenzhen Third People’s Hospital for the work they have done. This study was supported by the National Science Fund for Distinguished Young Scholars (82025022), the National Natural Science Foundation of China (82002140, 82171752, 82101861), the Guangdong Basic and Applied Basic Research Foundation (2021B1515020034, 2019A1515011197, 2021A1515011009, 2020A1515110656), and the Shenzhen Science and Technology Program (RCYX20200714114700046, JSGG20200207155251653, JSGG20200807171401008, KQTD20200909113758004, JCYJ20190809115617365).

## Author contributions

Z.Z. is the principal investigator of this study. Z.Z., B.J., and L.L. conceived and designed the study. B.J., B.Z., S.S., Q.F., and X.G. performed all experiments together with assistance from H.W., L.C., H.G., and D.S.. Z.Z. and B.J. wrote the manuscript and all authors read and approved this version of manuscript.

## Data availability statements

We are happy to share reagents and information in this study upon request.

## Conflict of interests

The authors have no conflict of interest.

---

**Figure S1.**
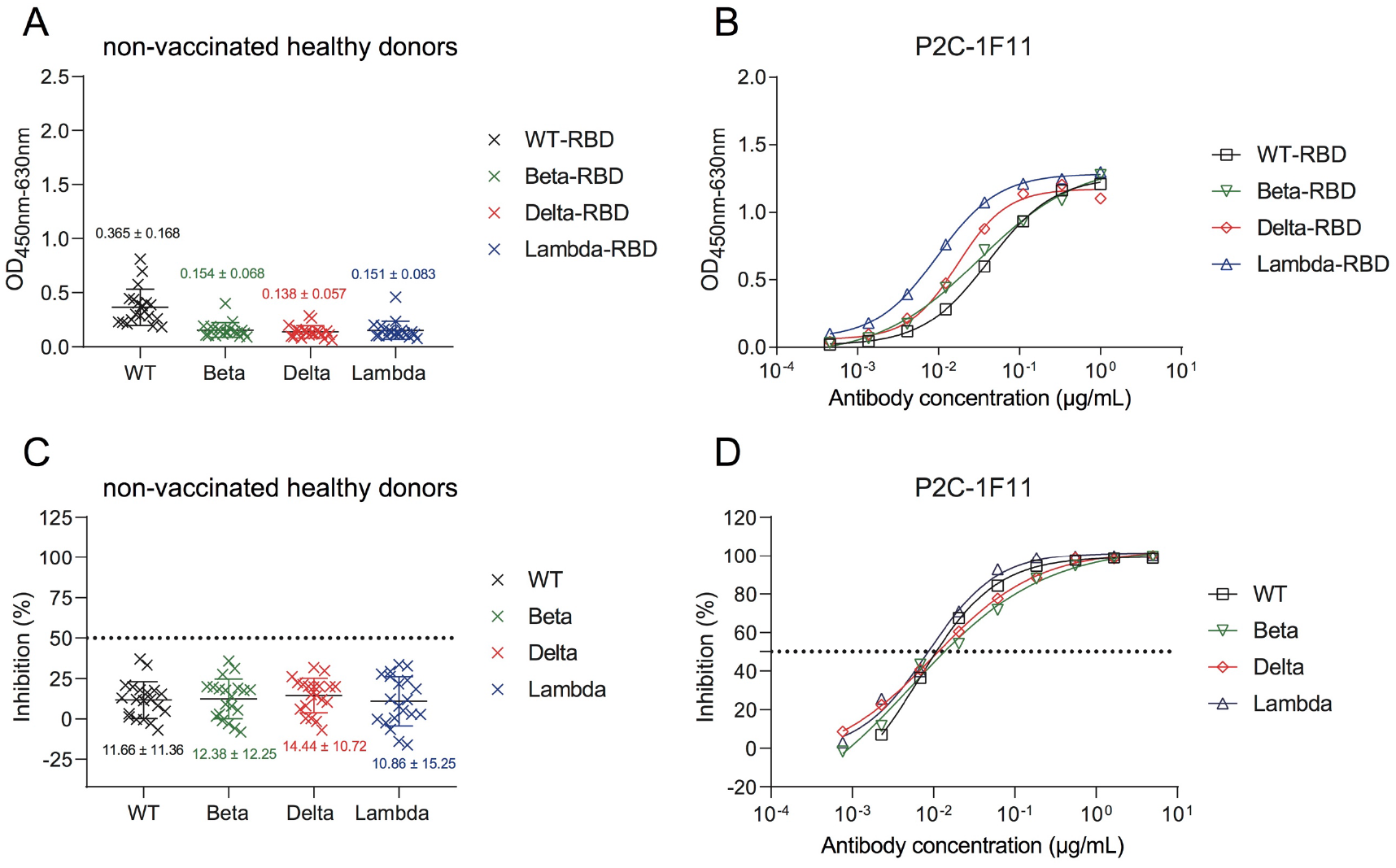
ELISA and neutralization profiles of non-vaccinated healthy donor plasma and a positive control mAb. ELISA binding of 20 non-vaccinated healthy donor plasma samples collected prior to COVID-19 pandemic **(A)** and a positive control mAb **(B)** to SARS-CoV-2 WT, Beta, Delta, and Lambda RBD proteins. Neutralizing activities of 20 non-vaccinated healthy donor plasma samples collected prior to COVID-19 pandemic **(C)** and a positive control mAb **(D)** against SARS-CoV-2 pseudoviruses of WT, Beta, Delta, and Lambda variants. Healthy donor plasma samples were tested at a dilution of 1:20. A positive control mAb (P2C-1F11) was serially 3-fold diluted from 1 μg/mL in ELISA and 5 μg/mL in neutralizing assay. All experiments were performed in duplicate and the mean ± SD values were shown.

**Figure S2.**
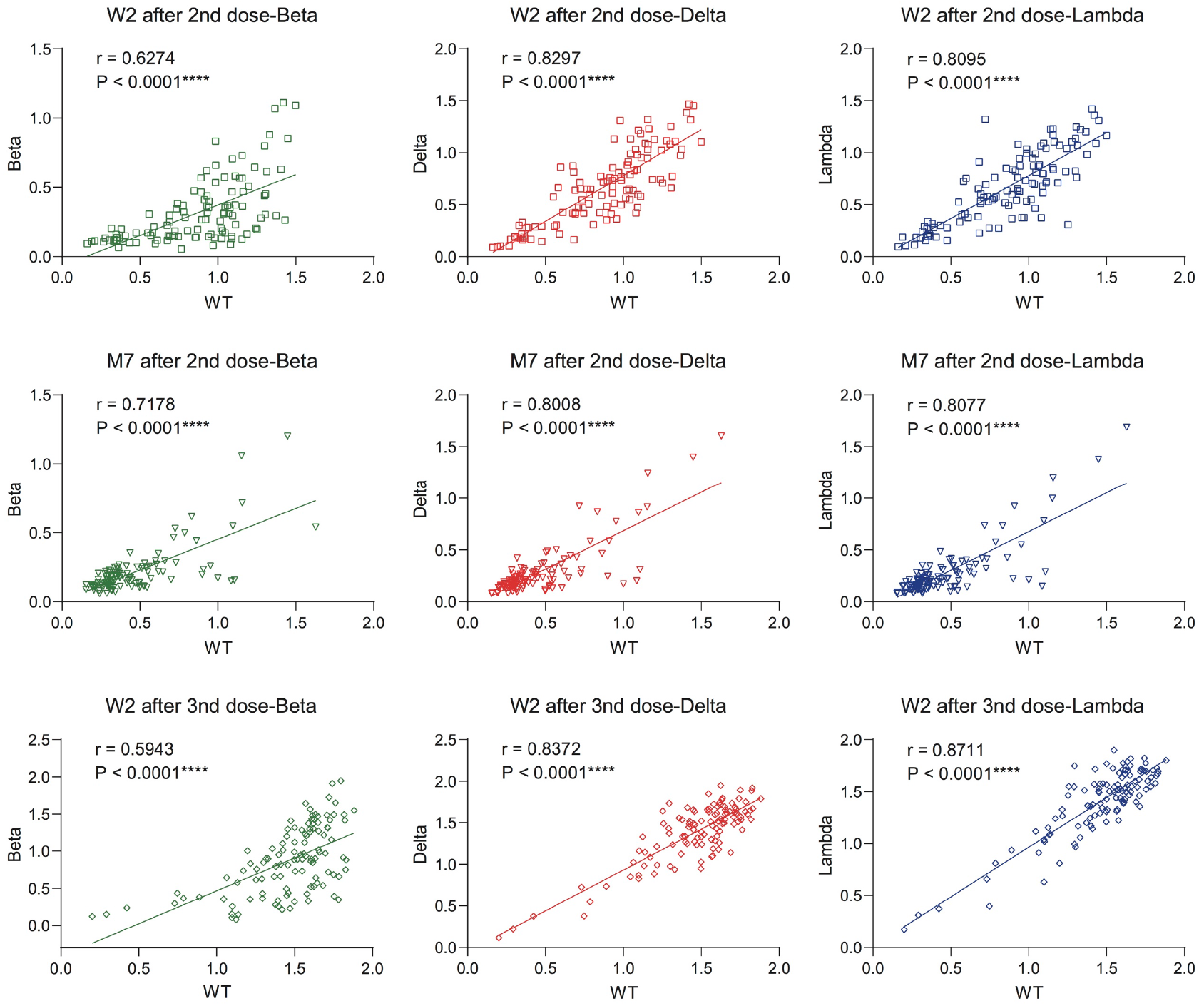
Correlation analysis between binding activities to SARS-CoV-2 WT and mutated (Beta, Delta, and Lambda) RBD proteins of 113 identical participants at three follow-up visits.

**Figure S3.**
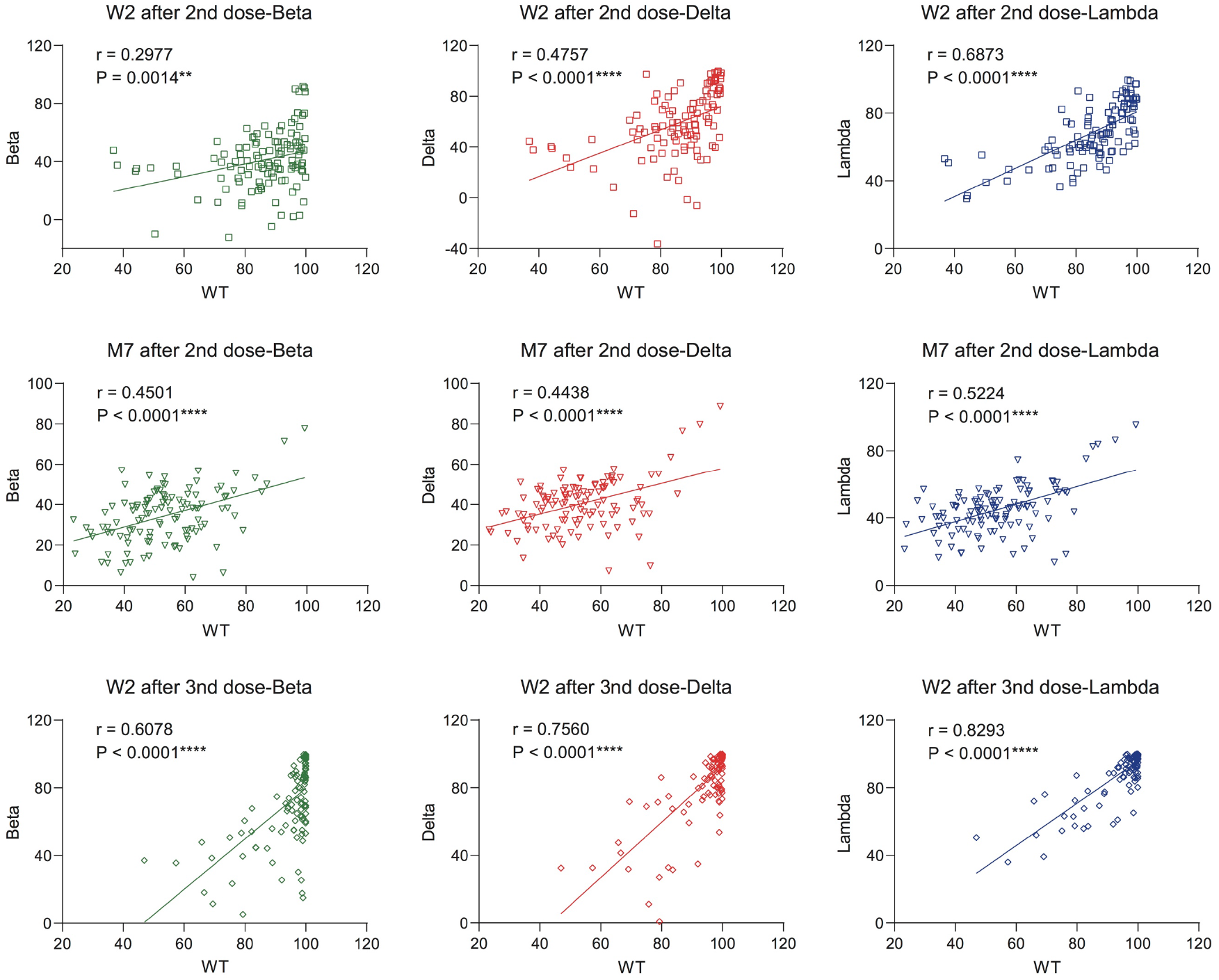
Correlation analysis between neutralizing activities against SARS-CoV-2 WT and variants (Beta, Delta, and Lambda) of 113 identical participants at three follow-up visits.

**Figure S4.**
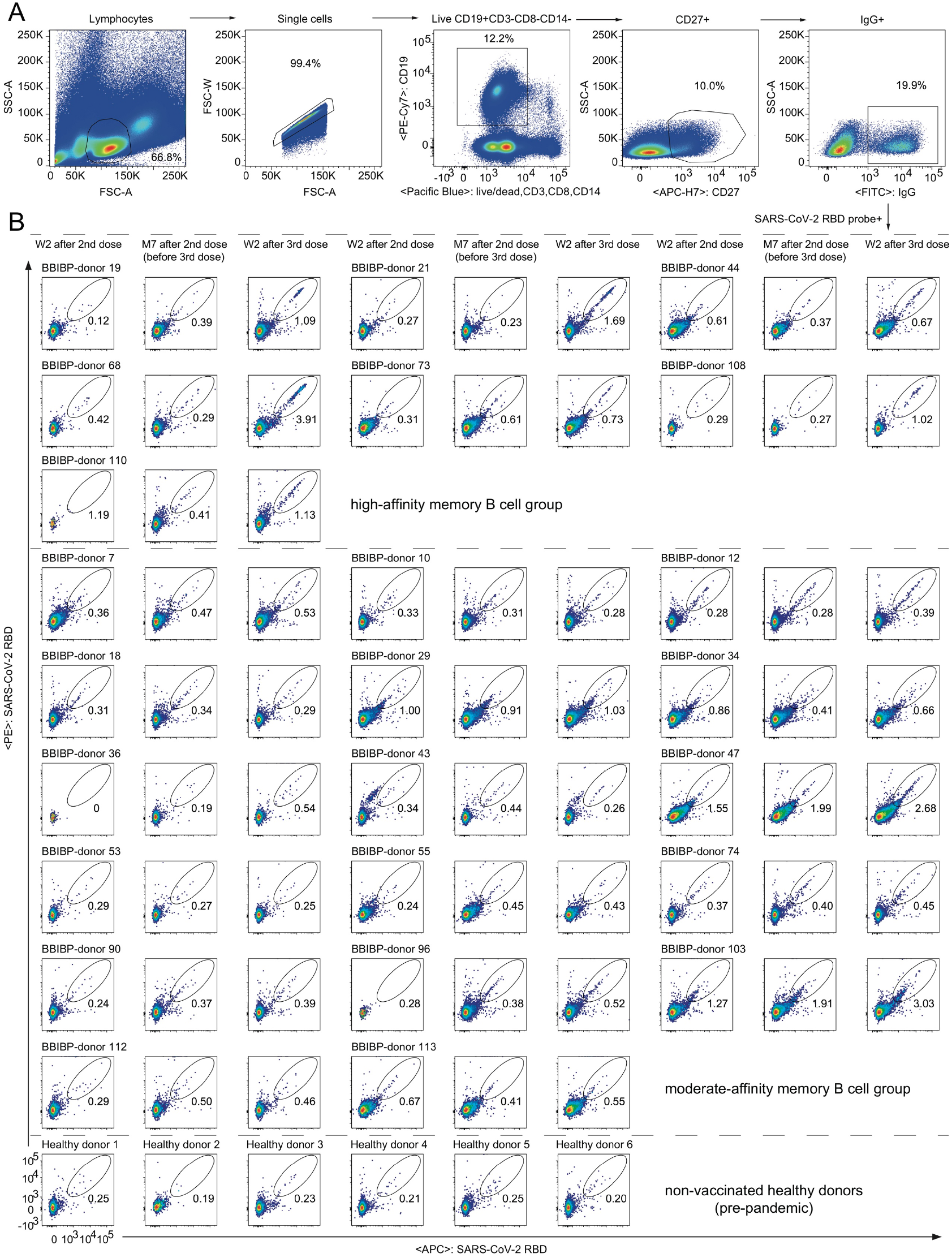
The gating strategy for identification of SARS-CoV-2 WT RBD-specific memory B cells by FACS. **(A)** Single B cells were gated as CD19^+^CD3^-^CD8^-^CD14^-^CD27^+^IgG^+^. **(B)** Flow cytometry showing the percentage of double-positive (APC^+^PE^+^) RBD-binding memory B cells of randomly selected 24 identical participants at three follow-up visits and 6 non-vaccinated healthy donors.

**Figure S5.**
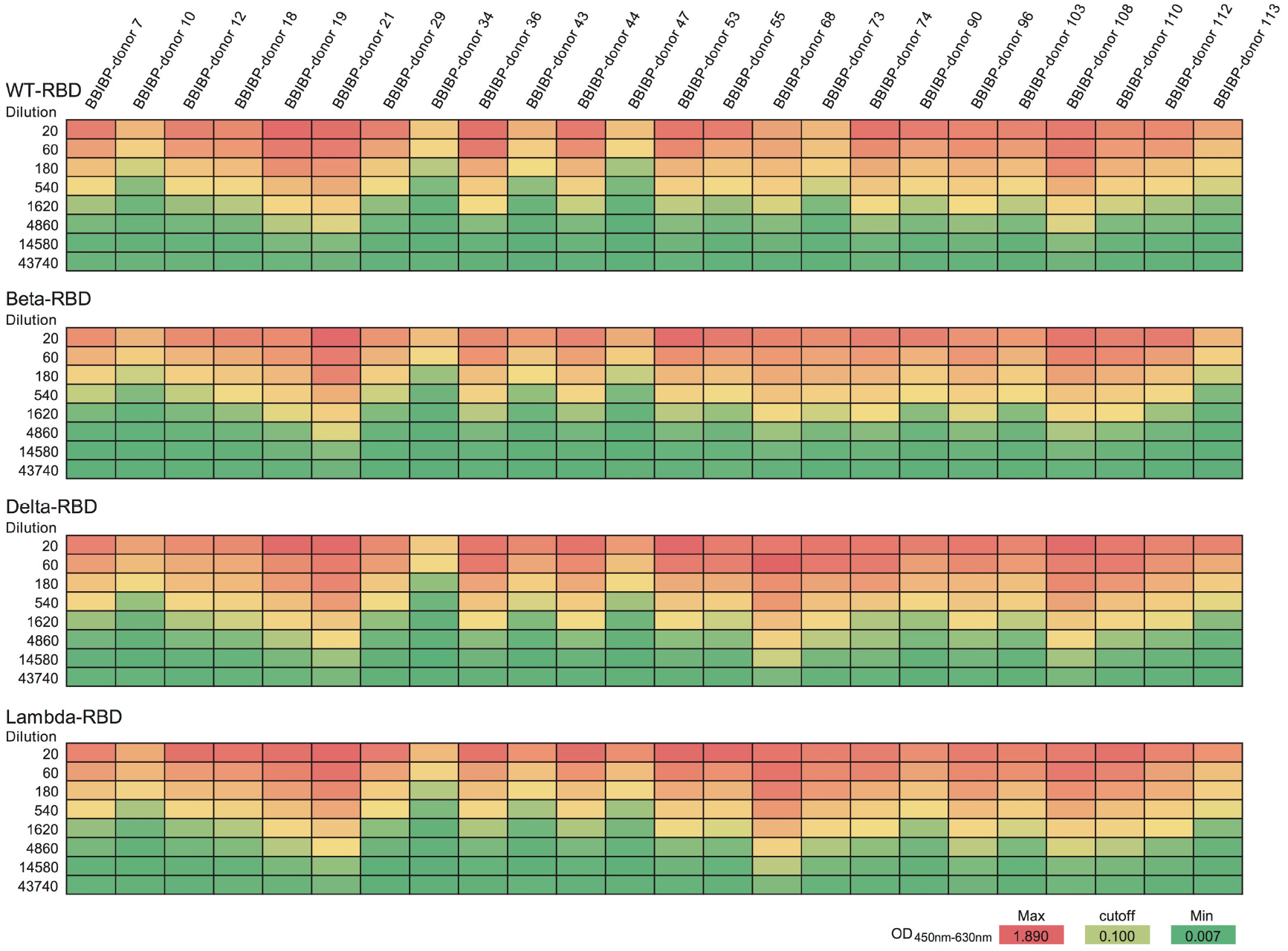
ELISA binding of 24 vaccinee plasma samples at Week 2 after third vaccination to SARS-CoV-2 WT and mutated (Beta, Delta, and Lambda) RBD proteins. All plasma samples were serially 3-fold diluted from 1:20. The assay was performed in duplicate and the mean value in each dilution was shown. The cut-off value was set as an OD_450nm-630nm_ value of 0.100 and the end-point titer was defined as the last dilution whose OD_450nm-630nm_ value was more than 0.100.

**Figure S6.**
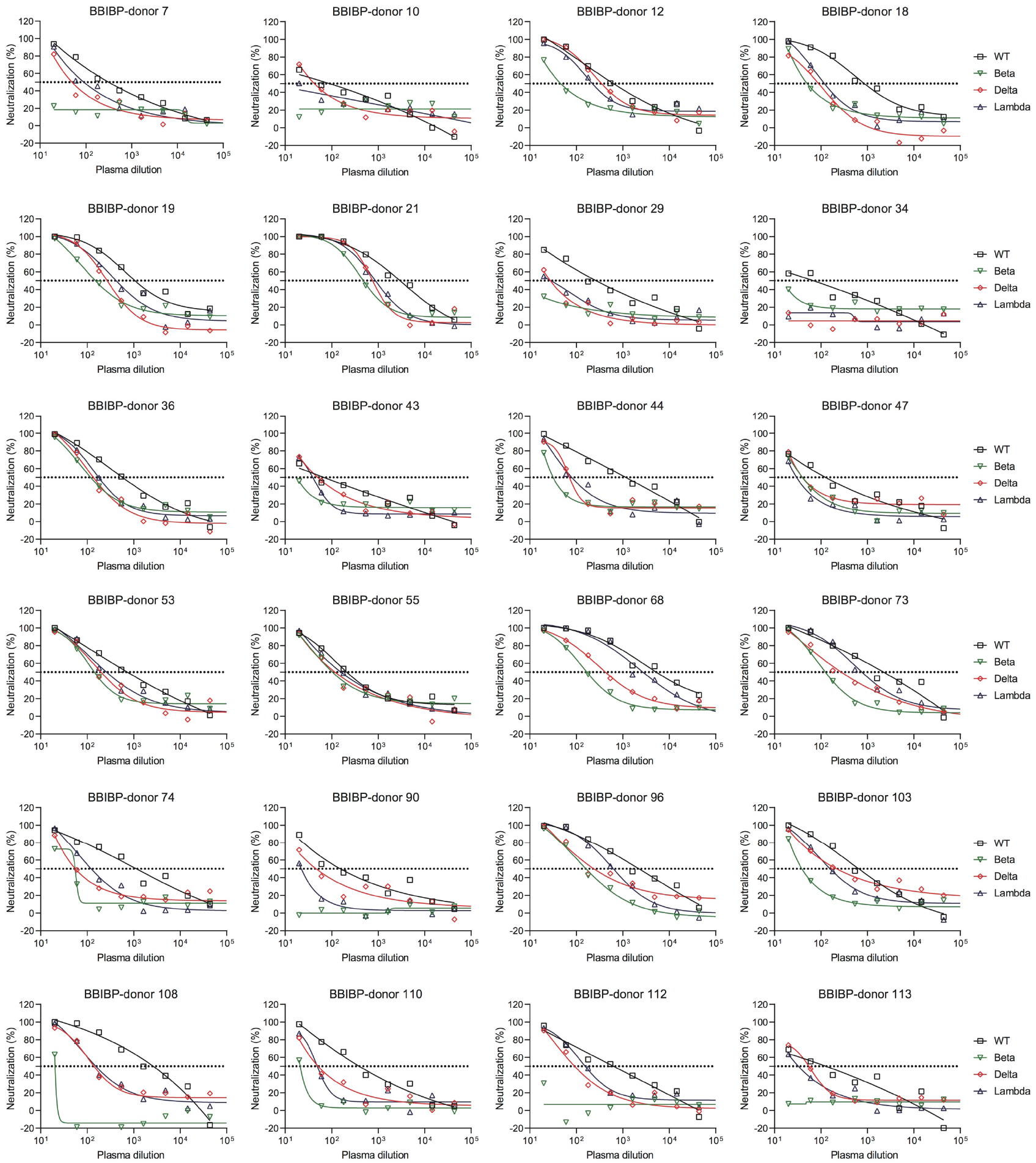
Neutralization curves of 24 vaccinee plasma samples at Week 2 after third vaccination against SARS-CoV-2 pseudoviruses of WT and variants (Beta, Delta, and Lambda). All plasma samples were serially 3-fold diluted from 1:20. The assay was performed in duplicate and the mean inhibition in each dilution was shown. A 50% reduction in viral infectivity was indicated by a horizontal dashed line.

## Notes

### Competing Interest Statement

The authors have declared no competing interest.

## References

1 Koff, W. C. et al. Development and deployment of COV D-19 vaccines for those most vulnerable. Sci Transl Med 13, doi:10.1126/scitranslmed.abd1525 (2021).

2 Dai, L. & Gao, G. F. Viral targets for vaccines against COV D-19. Nat Rev mmunol 21, 73–82, doi:10.1038/s41577-020-00480-0 (2021).

3 Subbarao, K. The success of SARS-CoV-2 vaccines and challenges ahead. Cell Host Microbe 29, 1111–1123, doi:10.1016/j.chom.2021.06.016 (2021).

4 Muik, A. et al. Neutralization of SARS-CoV-2 lineage B.1.1.7 pseudovirus by BNT162b2 vaccine-elicited human sera. Science, doi:10.1126/science.abg6105 (2021).

5 Wang, R. et al. Analysis of SARS-CoV-2 variant mutations reveals neutralization escape mechanisms and the ability to use ACE2 receptors from additional species. mmunity, doi:10.1016/j.immuni.2021.06.003 (2021).

6 Dejnirattisai, W. et al. Antibody evasion by the P.1 strain of SARS-CoV-2. Cell 184, 2939–2954 e2939, doi:10.1016/j.cell.2021.03.055 (2021).

7 Planas, D. et al. Reduced sensitivity of SARS-CoV-2 variant Delta to antibody neutralization. Nature, doi:10.1038/s41586-021-03777-9 (2021).

8 Liu, C. et al. Reduced neutralization of SARS-CoV-2 B.1.617 by vaccine and convalescent serum. Cell, doi:10.1016/j.cell.2021.06.020 (2021).

9 Kimura, . et al. SARS-CoV-2 Lambda variant exhibits higher infectivity and immune resistance. bioRxiv, doi:10.1101/2021.07.28.454085 (2021).

10 Kustin, T. et al. Evidence for increased breakthrough rates of SARS-CoV-2 variants of concern in BNT162b2-mRNA-vaccinated individuals. Nat Med 27, 1379–1384, doi:10.1038/s41591-021-01413-7 (2021).

11 Pollett, S. D. et al. The SARS-CoV-2 mRNA vaccine breakthrough infection phenotype includes significant symptoms, live virus shedding, and viral genetic diversity. Clin nfect Dis, doi:10.1093/cid/ciab543 (2021).

12 Hacisuleyman, E. et al. Vaccine Breakthrough nfections with SARS-CoV-2 Variants. N Engl J Med 384, 2212–2218, doi:10.1056/NEJMoa2105000 (2021).

13 Duerr, R. et al. Dominance of Alpha and ota variants in SARS-CoV-2 vaccine breakthrough infections in New York City. J Clin nvest 131, doi:10.1172/JC152702 (2021).

14 Falsey, A. R. et al. SARS-CoV-2 Neutralization with BNT162b2 Vaccine Dose 3. N Engl J Med, doi:10.1056/NEJMc2113468 (2021).

15 Xia, S. et al. Effect of an nactivated Vaccine Against SARS-CoV-2 on Safety and mmunogenicity Outcomes: nterim Analysis of 2 Randomized Clinical Trials. JAMA 324, 951–960, doi:10.1001/jama.2020.15543 (2020).

16 Xia, S. et al. Safety and immunogenicity of an inactivated SARS-CoV-2 vaccine, BB BP-CorV: a randomised, double-blind, placebo-controlled, phase 1/2 trial. Lancet nfect Dis 21, 39–51, doi:10.1016/S1473-3099(20)30831-8 (2021).

17 Barnes, C. O. et al. SARS-CoV-2 neutralizing antibody structures inform therapeutic strategies. Nature, doi:10.1038/s41586-020-2852-1 (2020).

18 McCallum, M. et al. SARS-CoV-2 immune evasion by the B.1.427/B.1.429 variant of concern. Science, doi:10.1126/science.abi7994 (2021).

19 Wang, P. et al. ncreased resistance of SARS-CoV-2 variant P.1 to antibody neutralization. Cell Host Microbe 29, 747–751 e744, doi:10.1016/j.chom.2021.04.007 (2021).

20 Lucas, C. et al. mpact of circulating SARS-CoV-2 variants on mRNA vaccine-induced immunity. Nature, doi:10.1038/s41586-021-04085-y (2021).

21 Liu, H. et al. The Lambda variant of SARS-CoV-2 has a better chance than the Delta variant to escape vaccines. bioRxiv, doi:10.1101/2021.08.25.457692 (2021).

22 Wang, P. et al. Antibody Resistance of SARS-CoV-2 Variants B.1.351 and B.1.1.7. Nature, doi:10.1038/s41586-021-03398-2 (2021).

23 Tortorici, M. A. et al. Broad sarbecovirus neutralization by a human monoclonal antibody. Nature 597, 103–108, doi:10.1038/s41586-021-03817-4 (2021).

24 Starr, T. N. et al. SARS-CoV-2 RBD antibodies that maximize breadth and resistance to escape. Nature 597, 97–102, doi:10.1038/s41586-021-03807-6 (2021).

25 Jette, C. A. et al. Broad cross-reactivity across sarbecoviruses exhibited by a subset of COV D-19 donor-derived neutralizing antibodies. Cell Rep 36, 109760, doi:10.1016/j.celrep.2021.109760 (2021).

26 Barros-Martins, J. et al. mmune responses against SARS-CoV-2 variants after heterologous and homologous ChAdOx1 nCoV-19/BNT162b2 vaccination. Nat Med 27, 1525–1529, doi:10.1038/s41591-021-01449-9 (2021).

27 Turner, J. S. et al. SARS-CoV-2 infection induces long-lived bone marrow plasma cells in humans. Nature 595, 421–425, doi:10.1038/s41586-021-03647-4 (2021).

28 Sokal, A. et al. mRNA vaccination of naive and COV D-19-recovered individuals elicits potent memory B cells that recognize SARS-CoV-2 variants. mmunity, doi:10.1016/j.immuni.2021.09.011 (2021).

